# From Farm to Pharmacy: Strawberry-Enabled Oral Delivery of Protein Drugs

**DOI:** 10.1101/2020.03.11.987461

**Authors:** Nicholas G. Lamson, Katherine C. Fein, John P. Gleeson, Sijie Xian, Alexandra Newby, Namit Chaudhary, Jilian R. Melamed, Kyle Cochran, Rebecca L. Ball, Kanika Suri, Vishal Ahuja, Anna Zhang, Adrian Berger, Dmytro Kolodieznyi, Brigitte F. Schmidt, Gloria L. Silva, Kathryn A. Whitehead

**Affiliations:** Department of Chemical Engineering, Carnegie Mellon University, Pittsburgh, Pennsylvania, United States; Department of Biomedical Engineering, Carnegie Mellon University, Pittsburgh, Pennsylvania, United States; Department of Chemistry, Carnegie Mellon University, Pittsburgh, Pennsylvania, United States

## Abstract

Although oral drug delivery is preferred by patients, it is not possible for proteins because the gastrointestinal tract is not sufficiently permeable. To enable the non-toxic oral uptake of protein drugs, we investigated plant-based foods as intestinal permeation enhancers, hypothesizing that compounds found in food would be well-tolerated by the gastrointestinal tract. Following a screen of over 100 fruits, vegetables, herbs, and fungi, we identified strawberry as a potent enhancer of macromolecular permeability in vitro and in mice. Natural product chemistry techniques identified pelargonidin, an anthocyanidin, as the active compound. In mice, insulin was orally administered with pelargonidin to induce sustained pharmacodynamic effects with doses as low as 1 U/kg and bioactivity of over 100% relative to the current gold standard of subcutaneous injection. Pelargonidin-induced permeability was reversible within two hours of treatment, and one month of daily dosing did not adversely affect mice as determined by weight tracking, serum concentrations of inflammatory markers, and tight junction gene expression. Results underscore the utility of plant-based foods in biomedical applications and demonstrate pelargonidin as an especially potent enhancer for the oral delivery of biologics.

## Body

Millions of patients are subjected daily to injections of single-dose therapeutics (e.g. vaccines) and chronic therapies (e.g. insulin). Unfortunately, the pain and inconvenience of injections leads to non-compliance. For example, 45%-60% of diabetic patients report that they have intentionally skipped doses of their medication as a result of injection-associated dread (*1, 2*). This non-compliance leads to poor disease control and long-term complications, which, in turn, exacerbate patient suffering and inflate healthcare costs (*3*). In contrast to injections, oral delivery is painless and convenient (*4*). Its development for protein and other macromolecular drugs would dramatically improve patient experience, compliance, and disease outcomes for a wide variety of maladies.

Unfortunately, oral protein delivery is not currently possible because the epithelial cell lining of the intestine inhibits the absorption of most macromolecules, especially protein drugs, into the bloodstream. Specifically, the epithelial cells are joined together by tight junction complexes that prevent the paracellular (between-the-cells) passage of any molecule larger than approximately 1 nm, or 600 Dalton (*5, 6*). Many delivery efforts have attempted to use chemical permeation enhancers to widen the diffusion channels in the tight junctions and achieve therapeutically significant absorption of oral macromolecules (*7, 8*) (**Fig 1a**). Unfortunately, most translational efforts have been thwarted by enhancer-caused cytotoxicity or structural damage to the small intestine (*9–11*), and oral delivery has been FDA-approved for only a handful of small peptide drugs (*12*).

**Figure 1:**
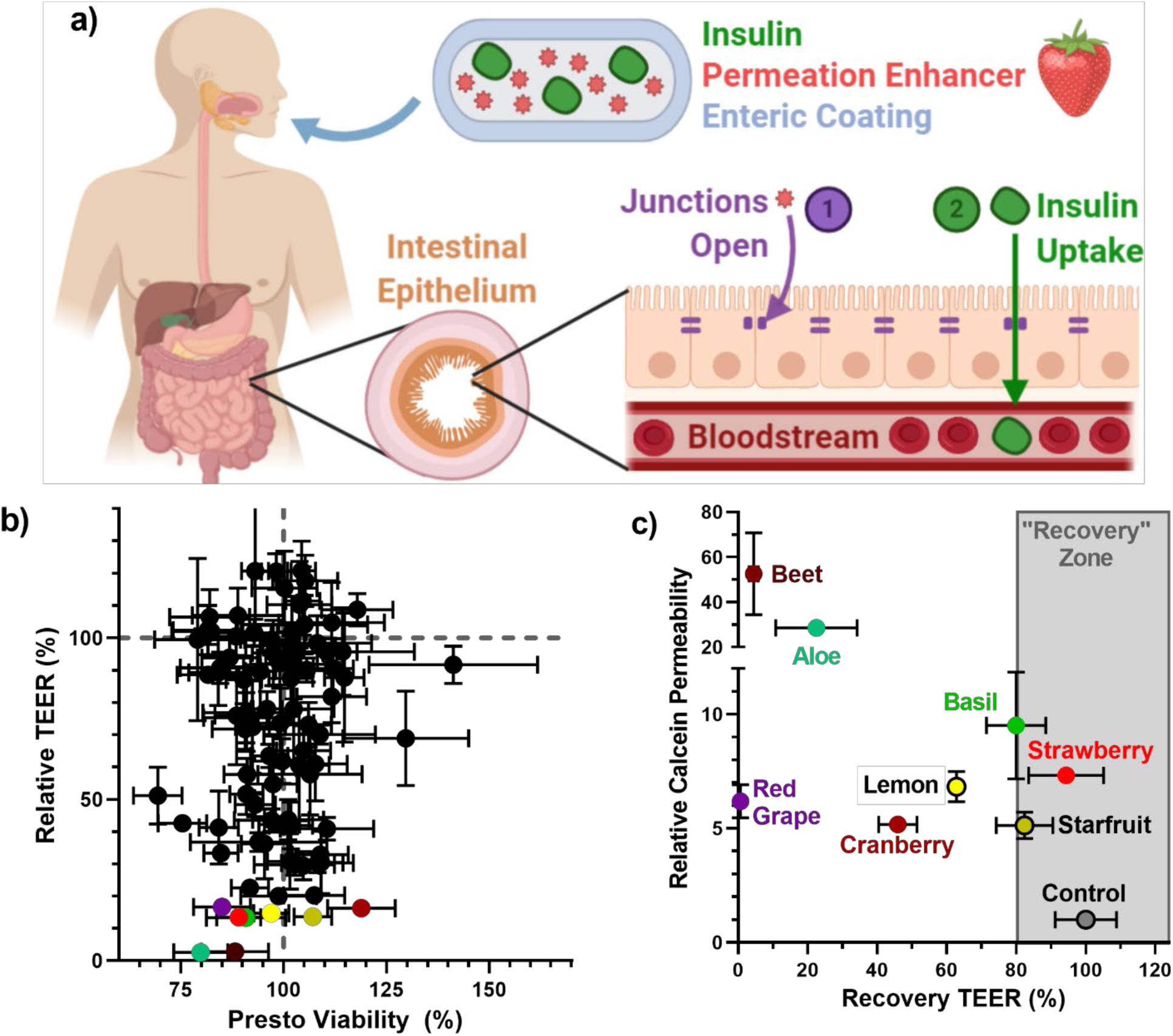
Plant-derived foods are a novel source of non-toxic permeation enhancers that enable oral protein delivery. **a)** A food-derived permeation enhancer is incorporated into a capsule containing a powdered protein drug (e.g. insulin) for oral administration. The capsule is coated with a pH-sensitive polymer (enteric coating) that delays release of the enhancer and drug until the capsule has localized to the small intestine. There, the enhancer opens the tight junctions between intestinal cells, enabling the protein drug to enter circulation. **b)** Food extracts varied in their effect on transepithelial electrical resistance (TEER, y-axis), which is a surrogate measurement for intestinal permeability. Lower TEER typically corresponds to higher drug permeability. **c)** The most effective TEER-reducing extracts increased the permeability of the model drug, calcein, across Caco-2 intestinal monolayers (y-axis). Strawberry was chosen as our top candidate because it enabled full recovery of TEER values within 24 hours of treatment removal (x-axis). Error bars display s.e.m. (n = 3 for TEER and calcein permeability, n = 4 for viability).

To address the urgent need for safe and effective intestinal permeation enhancers, we chose to investigate a library of materials that we hypothesized would be non-toxic in the gastrointestinal tract: plant-derived foods. Foods contain thousands of compounds that are well-tolerated by intestinal cells. Over millions of years, humans and plants have coevolved via mutualism: plants (especially fruits, vegetables, and herbs) provide a source of nutrition for humans, while humans disperse seeds through cultivation, harvest, preparation, and defecation (*13*). Given this cooperation, edible plants and their molecular building blocks typically do not induce toxic or immunogenic responses from the intestine. Furthermore, many plant-derived compounds (e.g. tannins, flavonoids) are beneficially bioactive, inducing anti-inflammatory, anti-microbial, or anti-cancer effects (*14–16*).

To determine the suitability of plant-derived foods as intestinal permeation enhancers, we assembled a library of 106 fruits, vegetables, and herbs (**Table S1)**, collected primarily from grocery stores and farmer’s markets. In addition, we grew some unusual fruits and vegetables, including otricoli orange berries and purple sweet potatoes, in the corresponding author’s backyard. Each food was prepared by washing, removing inedible portions (e.g. peels or pits), and blending with deionized water to form a slurry. The mash was then centrifuged and filtered to remove any water-insoluble materials, adjusted to neutral pH, and freeze-dried to yield a powdered extract. These extracts were dissolved at a concentration of 15 mg/mL in cell culture media for *in vitro* screening.

We began by examining the hypothesis that food-derived compounds are non-toxic to intestinal cells. Specifically, extracts were incubated with Caco-2 intestinal cells for three hours, then cell viability was measured using the PrestoBlue® assay (*17*) (**Table S1**). As expected, most fruit, vegetable, and herb extracts did not significantly reduce Caco-2 cell viability (**Fig S1, Fig 1b**). Those that did impact cell survival included garnishes and herbs that humans don’t typically consume in large or quantities (e.g. citrus rinds, rosemary). We also noted that several fruits known to contain protease enzymes (including kiwi, pineapple, and papaya) registered as cytotoxic according to the PrestoBlue® assay. Visual inspection revealed that these extracts were not killing the cells, but simply lifting them from the culture plate in the same manner that trypsin is used for dissociation during the cell culture passaging process.

We next took the non-cytotoxic extracts and examined their permeation enhancement efficacy by measuring the transepithelial electrical resistance (TEER) of treated Caco-2 monolayers (*17, 18*). TEER measures the resistance of the cell monolayers to ion passage, so a reduction in TEER indicates increased intestinal permeability (*19, 20*). Extracts produced a wide range of effects on permeability (**Fig 1b**). While many extracts did not impact TEER (y-axis values near the dashed gray line), a good number produced excellent permeation enhancement (TEER values less than 50%). Interestingly, a small number of extracts rendered the monolayers less permeable (e.g. raspberry, **Table S1**), which may be useful in the context of leaky gut therapy (*21*).

For the most promising permeation-enhancing extracts (colored points in **Fig. 1b**), we confirmed that TEER reductions corresponded with increased solute permeability. Specifically, we measured transport of the fluorescent molecule calcein, a 623 Da model drug, across Caco-2 monolayers (*17*) (y-axis, **Fig 1c**). As expected, all of these extracts increased calcein permeability. Improvements ranged from 5-fold (cranberry) to more than 50-fold (beet) compared to untreated control monolayers. Because reversibility of effect is important when selecting a permeation enhancer, we also measured the TEER values of intestinal monolayers 24 hours after extract removal. These data are presented as percentage recovery compared to untreated monolayers (x-axis, **Fig 1c**). We chose strawberry for future experiments because it offered the most promising combination of permeation enhancement and monolayer recovery.

Next, we focused on answering the most critical question for further technology development: Of the thousands of chemical compounds found in strawberry, which is responsible for strawberry’s unique ability to permeabilize the intestine reversibly and non-toxically? Upon review of our screening results, we noted an intriguing trend: some fruits and vegetables varied dramatically in their effect on intestinal permeability as a function of color. For example, red tomatoes, grapes, and potatoes were effective permeation enhancers. Yellow tomatoes, green grapes, and white potatoes, however, did not substantially affect monolayer permeability (**Fig 2a**). These data suggested that our active, permeation-enhancing compound contributes to plant coloration or is otherwise related to pigmented molecules.

**Figure 2:**
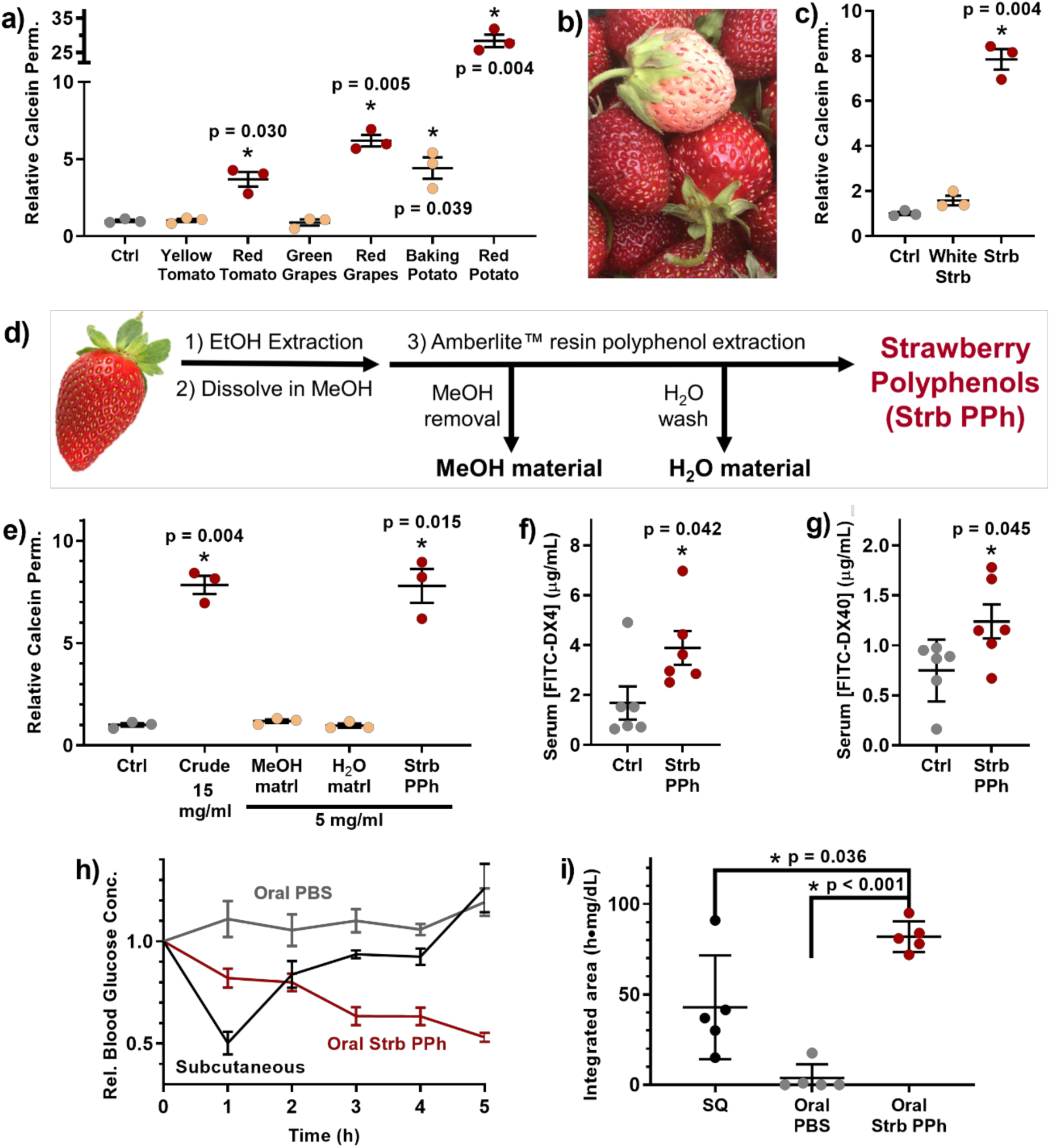
Polyphenolic compound(s) in strawberry enhance intestinal permeability. **(a)** Only the red varieties of several foods enhanced the permeability of Caco-2 monolayers. **(b)** White Carolina Pineberry strawberries were grown for this study. **(c)** White strawberries, which lack polyphenolic pigments, were not effective permeation enhancers. **(d)** Polyphenols were extracted from strawberry via adsorption to Amberlite™ resin and a sequence of washing steps. **(e)** Polyphenolic compounds isolated from strawberries enhanced intestinal permeability to the same extent as crude strawberry extract at one third of the dose. **(f)** Treatment with strawberry polyphenols doubled the uptake of orally-administered 4 kDa dextran (FITC-DX4) and **(g)** 40 kDa dextran (FITC-DX40) in mice. **(h)** 1 U/kg of insulin was delivered to mice by intestinal injection following oral delivery of either saline (control) or strawberry polyphenols. Subcutaneous injection of 1 U/kg insulin shown for comparison. Strawberry polyphenols induced sustained reductions in blood glucose levels. **(i)** Integrated areas above the curves from (h) demonstrate that strawberry polyphenols (Strb PPh) enabled significantly better insulin bioactivity than subcutaneous injection. Error bars display s.e.m. (n = 3 for panels (a), (c), and (e), n = 6 for panels (f-g), and n = 5 for panels (h-i)). * p < 0.05 w.r.t. control, unless otherwise denoted, by two-tailed t-test with Welch’s correction.

To test this theory, and because we like to eat home-grown strawberries, we grew a crop of the Carolina Pineberry, a white-fruited strawberry cultivar (**Fig 2b**). When applied to Caco-2 monolayers, these white strawberries failed to enhance intestinal permeability (**Fig. 2c**), confirming that we were searching for a compound that is related to strawberry coloration. The most abundant class of pigmented compounds in strawberries is the polyphenols (*22*), a chemically diverse family of molecules that each contain multiple phenol groups (*23*). Based on this connection, and the knowledge that white strawberries have multiple downregulated synthesis pathways for polyphenols (*24*), we proceeded to examine strawberry polyphenols as a narrowed set of potential permeation enhancers.

To isolate the polyphenols, we ground freeze-dried strawberries to a powder, extracted them with ethanol, and dried the resulting material. We then dissolved this in methanol and applied it to Amberlite™ XAD7 resin, which selectively adsorbs phenol groups (*17, 25*) (**Fig 2d**). Vacuum filtration of the resin removed all of the non-phenolic material with the methanol and subsequent water washes, and the polyphenols were eluted using ethanol. After drying, this separation process yielded approximately 3.5 grams of polyphenol extract per kilogram of strawberries (about 50 strawberries).

When applied to Caco-2 monolayers, the non-phenolic material did not affect permeability (**Fig 2e**). However, the polyphenols boosted calcein permeability to the same extent as the crude strawberry extract at only one third the concentration. With a purer permeation enhancer in hand, we next asked whether our cell culture results extended *in vivo*. For initial experiments, we orally dosed mice with strawberry polyphenols and 4 kDa FITC-labelled dextran (FITC-DX4), which is a non-digestible and fluorescent macromolecule commonly used to model oral uptake of small proteins (*17*). As expected, the strawberry polyphenols more than doubled FITC-DX4 absorption across the intestinal barrier (**Fig 2f**). We observed a similar increase in uptake for orally-administered 40 kDa FITC-dextran (FITC-DX40) (**Fig 2g**), suggesting that the strawberry polyphenols can be used to orally deliver mid-sized macromolecular drugs.

We then probed whether strawberry-derived polyphenols could enable the oral delivery of a functional protein. As a proof-of-concept, we chose to work with insulin (5.8 kDa) because 1) it is widely prescribed but not orally bioavailable, and 2) its bioactivity is readily assessed from a drop of whole blood. Specifically, successfully delivered insulin enables sugar uptake into cells and results in lower blood glucose levels. In these experiments, mice received an oral dose of strawberry polyphenols, followed by an injection of insulin directly into the small intestine to circumvent digestion in the stomach (*17*). Mice that received insulin and strawberry polyphenols experienced a substantial and sustained reduction in blood glucose concentration compared to mice that received insulin after a saline gavage (**Fig 2h**). Further, the strawberry polyphenols and intestinal insulin combination sustained hypoglycemia for at least three hours longer than the same 1 U/kg dose of subcutaneous insulin, the current gold standard of administration.

To compare the insulin bioactivity of these administration methods, we integrated the area between each mouse’s glucose curve and its starting blood sugar concentration. The areas above the curve (AACs) show that pharmacodynamic activity of intestinal insulin in polyphenol-treated mice is approximately double that of subcutaneous insulin (**Fig 2i**), yielding a relative bioactivity value of 191%. Additionally, the sustained activity of the polyphenol-assisted insulin indicates that the orally enhanced route may be advantageous for drugs that require extended release profiles.

Buoyed by these excellent proof-of-concept bioactivity data, we resumed our search for a single, active component of strawberry. To continue the compound isolation process, we turned to medium-pressure liquid chromatography (MPLC) for further resolution of the strawberry polyphenol extract (*17*) (**Fig S2**). Briefly, the full strawberry polyphenol mixture was separated once, and consecutive fractions were combined based on color and similarity of ultra performance liquid chromatography (UPLC) traces. The pooled samples were further separated by four additional runs of MPLC and re-combined based on UPLC traces. This yielded 22 fractions, including two pure compounds, for screening on Caco-2 monolayers. Although the vast majority of the samples did not affect epithelial permeability (**Fig 3a**), one fraction, ε3, was an exceptional permeation enhancer. The late elution of ε3 from the columns identified it as one of the less hydrophilic members of the polyphenol family. It was also deep red in color and a very small fraction, yielding less than three milligrams from one gram of polyphenol starting material. While the small quantity of product suggested that the fraction was particularly potent, it also complicated efforts to discern its molecular identity.

**Figure 3:**
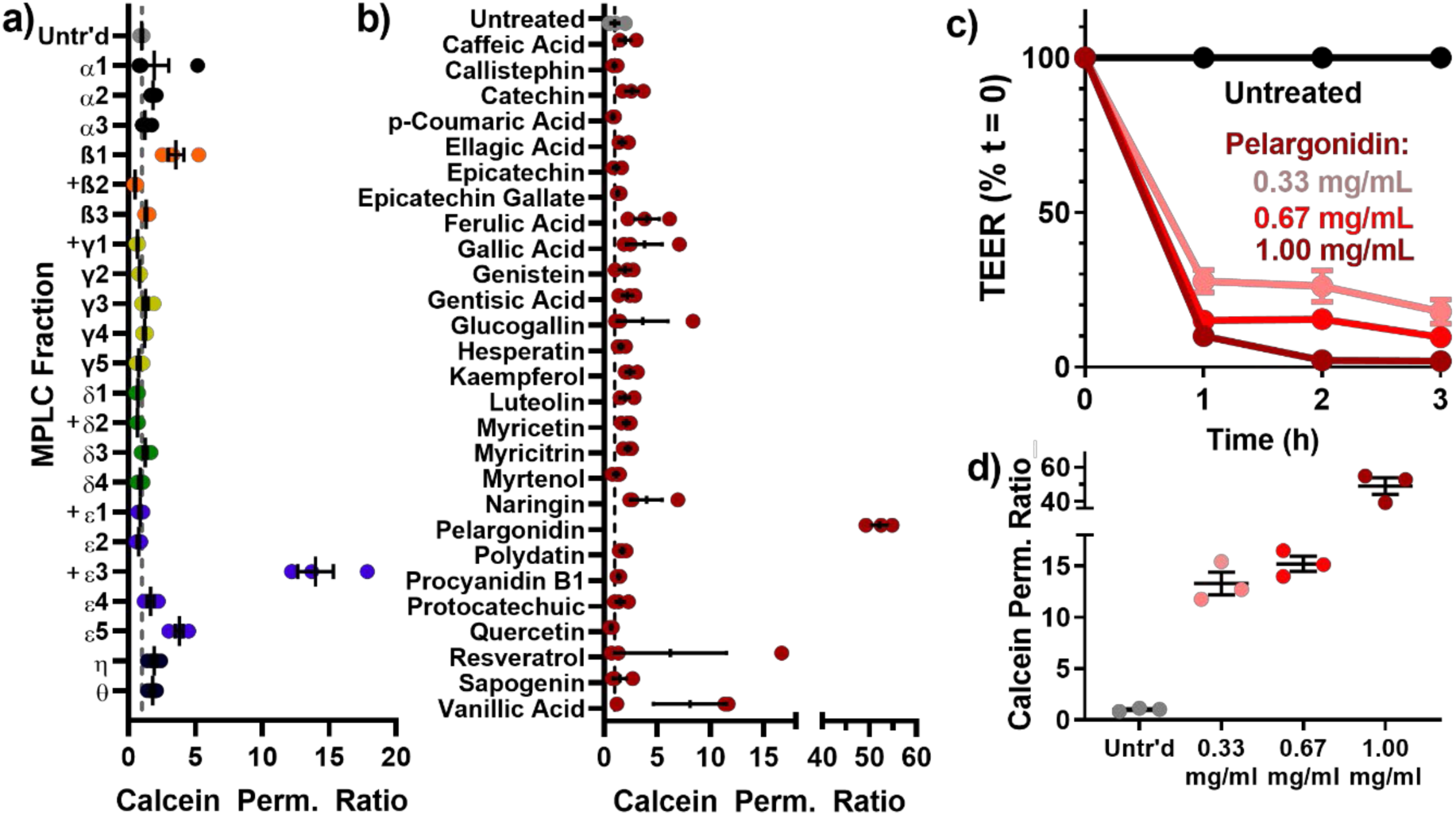
Chromatographic separation of strawberry polyphenols identified pelargonidin as the primary, active permeation enhancer. **(a)** Of 22 fractions that resulted from MPLC separations of strawberry polyphenols, only the ε3 fraction enhanced the permeability of Caco-2 monolayers. “+” denotes fractions that were too small to examine at the otherwise standardized concentration of 1 mg/mL. **(b)** Similarly, of a large group of commercially-purchased phenolic compounds known to occur in strawberries, only the pigment molecule pelargonidin significantly increased the permeability of calcein across cell monolayers. **(c)** Pelargonidin permeabilized intestinal Caco-2 cells, as measured by TEER, and **(d)** improved calcein permeability in a dose-dependent manner. Error bars represent s.e.m. (n = 3 - 4).

To expedite our identification of fraction ε3, we purchased a library of 27 known phenolic compounds from strawberry. As with the chromatography fractions, we screened these for their bioactivity on Caco-2 cells at 1 mg/mL concentration (**Fig 3b**). Fortuitously, one of these compounds was an effective permeation enhancer: pelargonidin. Pelargonidin and its glucoside, callistephin, contribute to the red coloration in strawberries (*26*), with the vast majority of the pigment is present in glycosylated form (*27*). However, our screening showed that callistephin was not an effective permeation enhancer, and that only the aglycone form of pelargonidin improved epithelial permeability. This is consistent with the observations that the effective fraction, ε3, yielded a small mass of deep red, fairly hydrophobic material. We further confirmed the common identity of ε3 and pelargonidin by mass spectrometry and nuclear magnetic resonance (NMR) (*17*) (**Figs S3** and **S4**). Treatment of Caco-2 monolayers with commercially-obtained pelargonidin led to dose-dependent reductions in TEER (**Fig 3c**) and improvements in calcein permeability (**Fig 3d**).

We then asked whether pelargonidin, like the strawberry polyphenol mixture, enables oral macromolecule delivery in mice. First, we examined the kinetics of permeation enhancement following oral pelargonidin treatment using FITC-DX4. Permeation enhancing effects were highest one hour after pelargonidin treatment (**Fig 4a)** and returned to PBS-treated control levels within an additional hour. These data suggest a swift reversibility of effect, assuaging a common concern that permeation enhancers permit the unregulated transport of toxins over prolonged periods. Experiments focusing on the size of the drug cargo showed that the boosts to uptake of orally-delivered 40, 70, and 150 kDa dextrans decreased with larger size (**Fig 4b**). This maintenance of size exclusion after treatment alleviates fears that pelargonidin-based permeation enhancement would allow for megadalton or larger material, such as virus capsids or bacteria, to migrate out of the intestines and into the body.

**Figure 4:**
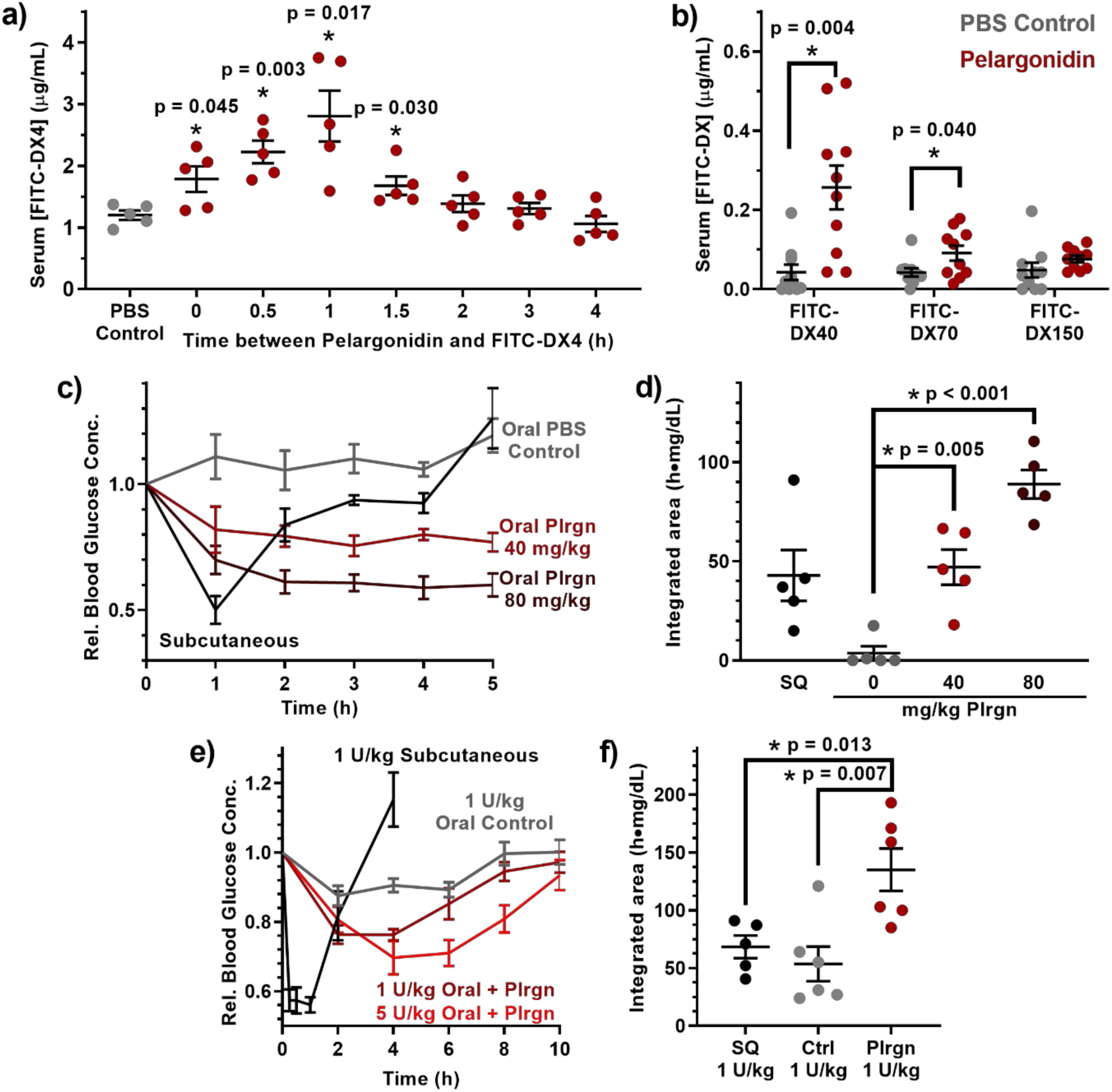
Pelargonidin, the active component of strawberry, enables oral delivery of functional proteins in mice. **(a)** Kinetic experiments of pelargonidin permeation enhancement in mice showed that the intestinal permeability 4 kDa FITC-dextran (FITC-DX4) peaks one hour after treatment and returns to baseline within another hour. **(b)** Pelargonidin treatment improved uptake of 40 kDa dextran, but increased permeability tapered off for larger dextrans. **(c)** The efficacy of intestinally injected insulin at a dose of 1 U/kg was dependent on the dose of oral pelargonidin pre-treatment, with **(d)** higher pelargonidin doses leading to higher bioactivity of the insulin. **(e**) Oral insulin doses of 1 U/kg (maroon) and 5 U/kg (red) reduced blood sugar in healthy mice when administered in pelargonidin-containing capsules. **(f)** Pelargonidin-insulin capsules resulted in double the bioactivity of subcutaneously injected insulin for 1 U/kg oral insulin. Error bars represent s.e.m. (n = 8-10 for panel b, n = 5-6 for all other experiments). *p < 0.050 w.r.t. control, unless otherwise denoted, by two-tailed t-test with Welch’s correction.

We again used insulin to demonstrate that pelargonidin enables the delivery of functional protein drugs across the intestinal barrier. Following oral pelargonidin administration, intestinal injections of insulin (1 U/kg) induced sustained decreases in blood glucose over at least four hours, while a subcutaneous injection induced a pronounced but brief response (**Fig 4c**). Area above the curve calculations showing that pharmacodynamic activity of intestinal insulin at the highest pelargonidin dose (80 mg/kg) is approximately double that of subcutaneous insulin (**Fig 4d** and **Table 1**). This performance outpaces other promising, recently reported strategies for oral insulin delivery, including ionic liquid (*28*) and anionic nanoparticles (*29*), which achieve between 60% and 100% relative bioactivity.

**Table 1:** Summary of relative bioactivity values for insulin delivered with or without pelargonidin absorption enhancer in healthy mice. Data is presented as arithmetic average ± standard error. Plrgn = pelargonidin. rBGmin = minimum average relative blood sugar achieved. AAC = insulin dose adjusted area above the blood glucose curve. rBA = dose-adjusted relative bioactivity. SQ = subcutaneous injection.

| Delivery Route | Insulin Dose | Pelargonidin | rBGmin | AAC | rBA |
| --- | --- | --- | --- | --- | --- |
| | $\frac{U}{kg}$ | $\frac{mg}{kg}$ | % | $\frac{h \cdot mg/dL}{U/kg}$ | % |
| SQ (5 h) | 1 | 0 | 48.3 | 42.9 | 100.0 |
| Intestinal (5 h) | 1 | 0 | 100.0 | 3.7 | 8.7 |
|  | 1 | 40 | 75.5 | 47.1 | 109.7 |
|  | 1 | 80 | 58.9 | 88.9 | 207.2 |
| SQ (10 h) | 1 | 0 | 56.1 | 68.5 | 100.0 |
| Oral (10 h) | 1 | 0 | 87.6 | 53.4 | (-9.8) |
|  | 1 | 40 | 76.3 | 135.2 | 140.2 |
|  | 5 | 40 | 69.7 | 152.0 | 34.2 |
|  | 0 | 40 | 86.4 | 59.0 | 0.0 |

Next, to demonstrate the efficacy of a fully-oral delivery system, we loaded pelargonidin and insulin together into mouse-sized capsules and coated them with the enteric polymer Eudragit® L100-55 to protect them from acid- and enzyme-mediated degradation in the stomach (*17*). When orally administered to mice, an insulin dose of only 1 U/kg decreased blood glucose concentrations for over eight hours compared to the same dose delivered without pelargonidin. In contrast, a 1 U/kg subcutaneous injection of insulin produced a sharp hypoglycemic effect that resolved within three hours (**Fig 4e**). Area above the curve calculations determined that the orally administered insulin was twice as bioactive as subcutaneous insulin for the 1 U/kg capsules (**Fig 4f**). To our knowledge, this is the highest oral insulin bioactivity reported in the literature. Mucoadhesive intestinal patches (*30*) and anionic nanoparticle permeation enhancers (*29*) in healthy animals have achieved 7% and 29% relative insulin bioactivity, whereas this system produces 140% of the pharmacodynamic potency per insulin molecule (**Table 1**). It is also notable that the hypoglycemic effect here is achieved using only a 1 U/kg oral insulin dose, as opposed to the 10-100 U/kg doses that other oral delivery systems have necessitated (*28–30*).

Having demonstrated the unusual efficacy of pelargonidin as an intestinal permeation enhancer for oral protein delivery, we conducted additional mechanistic and toxicity experiments to evaluate its potential for long-term use. First, we examined the rearrangement of tight junction proteins upon pelargonidin treatment in Caco-2 cells via immunofluorescence and confocal microscopy (*17*). While nuclear staining and mid-cell actin were consistent between untreated (**Fig 5a**) and pelargonidin-treated cells (**Fig 5b**), pelargonidin caused a marked rearrangement in actin at the apical (luminal) cell surface (**Figs 5c-d**), which is in the vicinity of tight junctions (*31*). The junction anchoring protein, zonula occludens 1 (ZO-1) (**Figs 5e-f**), and the tight junction protein, occludin (**Figs 5g-h**), delocalized from the tight junctions and displayed large gaps in the normally continuous rings at the boundary of each cell. By pre-treating Caco-2 monolayers with a collection of small molecule enzyme inhibitors, we determined that myosin light chain kinase (MLCK), an enzyme known to remodel the actin cell skeleton (*32*), is a critical mediator of pelargonidin-induced permeability (**Fig S5**). Inhibition of rho-associated protein kinase (ROCK) or kinase c-Src diminished but did not fully prevent the efficacy of pelargonidin. None of the other signaling enzymes that we examined proved necessary to the permeabilization process.

**Figure 5:**
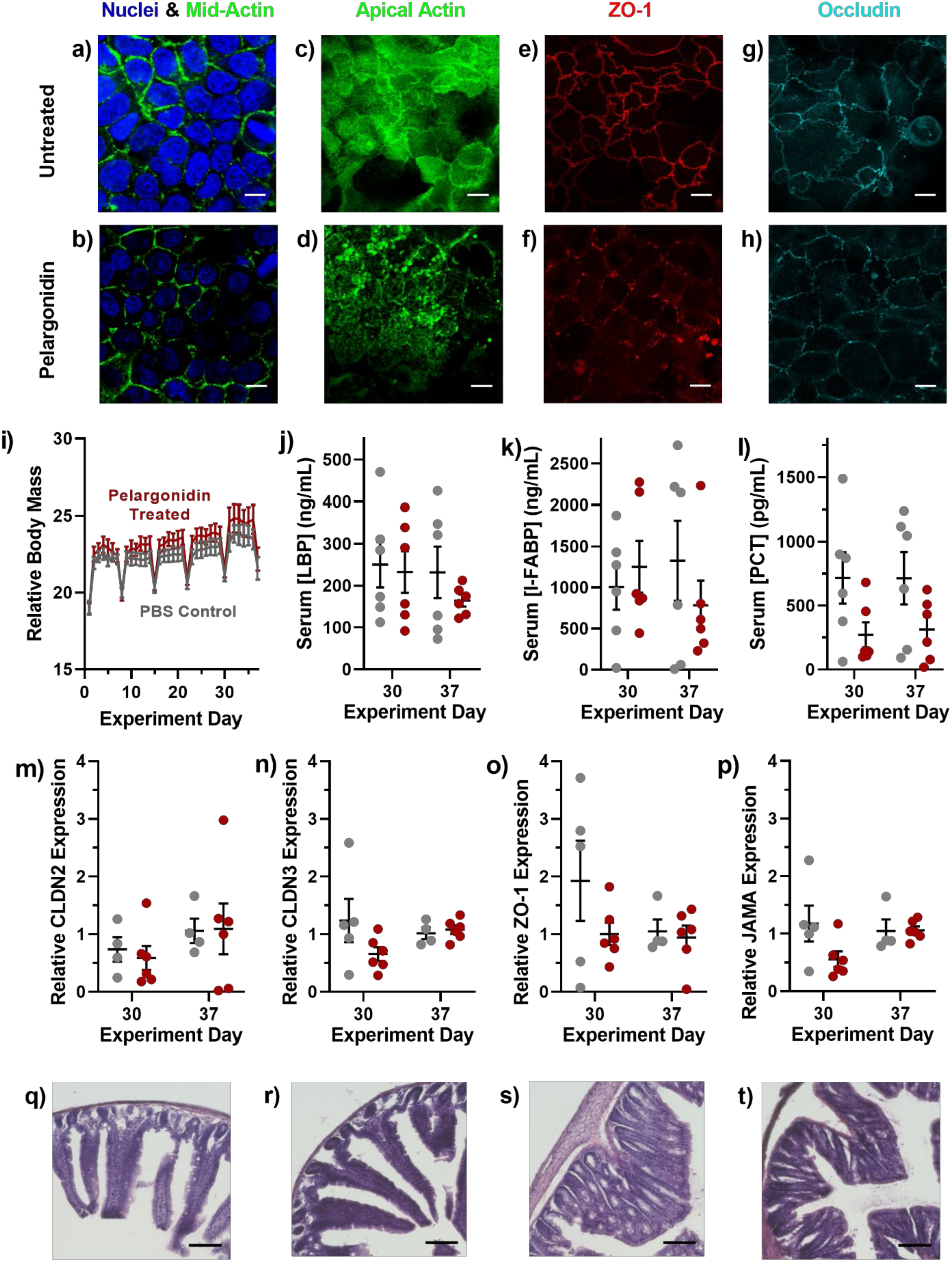
Pelargonidin induces reversible, non-toxic opening of intestinal tight junctions. **(a)** Compared to untreated Caco-2 cells, **(b)** pelargonidin-treated monolayers showed no difference in morphology for nuclei or mid-cell actin. **(c, d)** However, actin at the apical surface, as well as the tight junction proteins **(e, f)** ZO-1 and **(g, h)** occludin rearranged into more punctate forms as a result of pelargonidin treatment. **(i)** During one month of daily pelargonidin treatment, mice did not lose weight compared to control animals. Periodic weight loss was observed in both control and treated groups as a result of overnight fasting for weekly checkups. **(j)** Treated mice did not develop elevated levels of the inflammation markers lipopolysaccharide binding protein (LBP), **(k)** intestinal fatty acid binding protein (I-FABP), or **(l)** procalcitonin (PCT). qRT-PCR revealed no statistical difference in mRNA expression of the tight junction proteins **(m)** Claudin 2, **(n)** Claudin 3, **(o)** ZO-1, or **(p)** JAMA in the small intestines of control and pelargonidin-treated mice. **(q)** Representative histological images of small intestines from control mice and **(r)** pelargonidin treated mice displayed no tissue damage resulting from treatment. **(s)** There were also no discernable histological differences between control and **(t)** treated mouse colons. White scale bars are 10 µm and black scale bars are 100 µm. Error bars represent s.e.m. (n = 4 - 12). For panels i through p, no comparisons between treated and control mice achieved statistical significance by two-tailed t-test with Welch’s correction.

To address concerns that pelargonidin permeation enhancing treatment may cause damage to intestinal mucosae (*11*), we performed a long-term safety experiment. Specifically, we orally administered a 40 mg/kg pelargonidin dose to mice every day for thirty days. Half of the animals were further monitored for an additional, seven-day “recovery” period after the treatment regimen. In each case, we compared several intestinal and systemic health outcomes in pelargonidin-treated subjects to control animals receiving saline gavages (*17*). Treated mice did not lose weight during the study (**Fig 5i**) nor did they experience elevated blood levels of the intestinal inflammation markers lipopolysaccharide binding protein (*33*) (LBP, **Fig 5j**), intestinal fatty acid binding protein (*34, 35*) (I-FABP, **Fig 5k**), or procalcitonin (*35*) (PCT, **Fig 5l**). Further, gene expression analysis of mice’s small intestinal epithelia revealed no significant changes in the expression of the pore-forming tight junction protein Claudin 2 (*8*) (CLDN2, **Fig 5m**) or the barrier-forming proteins Claudin 3 (CLDN3, **Fig 5n**), Zonula Occludens 1 (ZO-1, **Fig 5o**) and Junctional Adhesion Molecule A (JAMA, **Fig 5p**) (*36*).

Finally, we collected intestinal samples from treated mice and subjected them to hematoxylin and eosin staining for histological analysis (*17*). Comparison of small intestines between control mice (**Fig 5q**) and pelargonidin treated mice (**Fig 5r**) revealed no immune cell invasion, necrosis, or discernible changes in tissue architecture. Similarly, there were no observable differences between colon tissue from control (**Fig 5s**) and pelargonidin treated (**Fig 5t**) animals. Taken together, these data indicate that daily intake of pelargonidin, at doses effective for oral drug delivery, is unlikely to cause short or long term intestinal or systemic toxicity.

We have shown here that pelargonidin, an anthocyanidin molecule abundant in strawberries, reversibly opens intestinal tight junctions, allowing uptake of orally-administered protein drugs into the blood stream. Further, the unusual absorption enhancing efficacy achieved with this compound suggests that it could provide sufficient bioactivity of therapeutic proteins to replace subcutaneous injections, broadly improving patient experience, compliance, and clinical outcomes for a wide variety of treatments and diseases.

## Acknowledgments

The authors thank Lynn Walker for her advice and for supplying ripe quinces from her home garden.

## Author contributions

Conceptualization, N.G.L., K.C.F., J.P.G., and K.A.W.; Methodology, N.G.L., K.C.F., J.P.G., R.L.B., and A.B.; Investigation, N.G.L., K.C.F., J.P.G., S.X., A.N., N.C., J.R.M., K.C., R.L.B., K.S., V.A., A.Z., A.B., D.K., and B.F.S; Resources, B.F.S.; Writing – Original Draft, N.G.L; Writing – Reviewing and Editing, K.C.F, J.P.G., A.N.N., and K.A.W.; Visualization, N.G.L., K.C.F., and K.A.W.; Supervision, K.A.W; Project Administration, N.G.L., K.C.F., B.F.S., G.L.S., and K.A.W.; Funding Acquisition, N.G.L. and K.A.W.

## Data and materials availability

All data are available in the manuscript or the supplementary materials.

## Supplementary Materials

Materials and Methods

Figs S1 – S6

Tables S1 – S3

