## Supplementary Information for "From Farm to Pharmacy: Strawberry-Enabled Oral Delivery of Protein Drugs"

**This file includes:**

Materials and Methods

Figures S1 to S6

Tables S1 to S3

Materials and Methods

Materials
 Penicillin/streptomycin, trypsin-ethylenediaminetetraacetic acid (trypsin-EDTA), phosphate buffer saline (PBS), fetal bovine serum (FBS), rat tail Collagen I, PrestoBlue® viability kits, Hoechst 33342, calcein, DAPI, AlexaFluor 488® conjugated phalloidin, AlexaFluor® 594 conjugated Anti-ZO-1 antibodies, and AlexaFluor® 594 Anti-Occludin antibodies were purchased from Life Technologies® (Thermo Fisher subsidiary, Carlsbad, CA, USA). Caco-2 cells were purchased from American Type Culture Collection® (ATCC, Manassas, VA, USA). Dulbecco’s Modified Eagles Medium (DMEM), Amphotericin B, Falcon® 225 cm^2^ tissue culture flasks, Falcon® HTS 24-Multiwell Insert Systems with 1µm pores, Falcon® 24-well plates, Corning® CellBIND® 96-well microplates, sodium butyrate, MITO+ serum extender, Aimstrip® Plus blood glucose strips, blood glucose monitor, gentisic acid, furoic acid, ellagic acid, kaempferol, naringin, vanillic acid, protocatechuic acid, ferulic acid, caffeic acid, lactone hexose, resveratrol, luteolin, bovine serum albumin (BSA), and OCT mounting media were obtained from VWR® (Radnor, PA, USA). FITC-labelled dextrans, Amberlite™ XAD7 resin, recombinant human insulin, aprotinin, catechin, epicatechin, polydatin, myricitrin, myricetin, hesperetin, myrtenol, gallic acid, metoclopramide hydrochloride, streptozotocin (STZ), U-73122, SR 3677, PP2, API-1, AZ 628, PIK, Harris modified hematoxylin, and alcoholic eosin Y were purchased from Sigma-Aldrich® (St. Louis, MO, USA). C18 bulk silica gel SMT-Bod-C18 was purchased from Separation Methods Technologies (Newark, DE, USA). Callistephin and procyanidin B1 were obtained from Alkemist Labs (Costa Mesa, CA, USA). Epicatechin gallate and pelargonidin were from ChromaDex (Irvine, CA, USA). Genistein, glucogallin, and sarsasapogenin were purchased from Toronto Research Chemicals (North York, ON, Canada). Mouse-sized (M) capsules and dosing kit were supplied by Torpac and Eudragit L100-55 enteric coating polymers were a gift from Evonik. DMSO-d_6_ and TMS were purchased from Cambridge Isotope Laboratories (Tewksbury, MA). The cDNA reverse transcriptase kit; primers for Beta-Actin, ZO-1, Claudin-2, Claudin-3, and JAM-A; and SYBR Select Master Mix were ordered from Applied Biosystems (Foster City, CA). ELISA kits for procalcitonin (PCT), lipopolysaccharide binding protein (LBP) and fatty acid binding protein 2 (I-FABP) were purchased from Abclonal (Woburn, MA).

Preparation of Crude Food Extract Library
Food samples were obtained from local supermarkets and farmer’s markets or grown from nursery seed/stock (Baker Creek Heirloom Seeds, Stark Bro’s Nurseries and Orchards Co.) in KAW’s garden, totaling 106 fruits, vegetables, and herbs (**Table S1**). After removing inedible portions (e.g. seeds and stems), the samples were blended with 125% w/w distilled water on medium-high speed for three minutes using a household blender. The resulting slurry was transferred to 50 mL conical tubes and centrifuged for 30 minutes at 400 RCF and room temperature. The resulting liquid was centrifuged to remove insoluble components, and the supernatant filtered through standard coffee filters to remove any large particulate matter, isolating water-soluble components that are compatible with aqueous cell culture assays. The extracts were then adjusted to neutral pH (7) with 1 M NaOH, and lyophilized. The resulting powders were stored at -80°C until use, when they were dissolved at 15 mg/mL in cell culture media immediately before testing.

Cell Culture
 Caco-2 lines were confirmed mycoplasma free by direct DNA staining with Hoechst 3334229. Cells were cultured in DMEM supplemented with 10% FBS, 100 IU/mL of penicillin, 0.1 mg/mL streptomycin, and 0.25 μg/mL Amphotericin B (“Caco-2 media”). Cultures were incubated at 37°C in a fully humid, 5% CO_2_ environment. The cells were subcultured with 0.25% trypsin-EDTA and subsequent passaging every 3 to 4 days at ratios between 1:3 and 1:8. Cells at passage numbers 20–50 were utilized for further experiments.

PrestoBlue® Assay
Caco-2 cells were seeded in a clear-bottom, black, 96-well plate at a concentration of 10^4^ cells/well. After incubating the plate overnight at 37°C, the media in the wells was aspirated and replaced with the treatment solutions (15 mg/mL, 100 μL/well). After three hours of exposure, the extracts were aspirated. PrestoBlue® reagent (10 μL/well) and Caco-2 media (90 μL/well) were added to the wells. Thirty minutes later, a BioTek® Synergy2 automated plate reader was used to measure the fluorescent signal produced by viable cells. The viability of each treatment is expressed as the ratio of the fluorescence intensity of the untreated cells to that of the untreated cells.

Caco-2 Permeability Experiments
 For transepithelial electrical resistance (TEER) and diffusion marker permeability experiments, TRIM models of rapid, 3-day Caco-2 intestinal epithelial monolayers were employed (*1*). Briefly, Caco-2 cells were suspended in DMEM supplemented with MITO+ serum extender (basal seeding medium, BSM), seeded at a density of 2×10^5^ cells per well on collagen-coated transwell membrane supports, and incubated for 24-48 hours. The media was then changed to DMEM supplemented with MITO+ and 2 mM sodium butyrate (enterocyte differentiation medium, EDM), and incubated for 48 hours. The TEER was monitored to confirm proper barrier formation, and only monolayers with initial TEER values of at least 150 Ω·cm^2^ were utilized for TEER or molecular permeability experiments.

Transwell inserts containing Caco-2 monolayers were transferred to 24-well plates containing 1 mL DMEM per well and allowed to equilibrate for 30 minutes before recording initial resistance values using a Millicell® voltohmmeter. Treatments were suspended in EDM (15 mg/mL unless otherwise specified) and applied to the apical chambers, and negative control wells received fresh EDM. TEER readings were taken after 15, 30, 60, 120, and 180 minutes. After 180 minutes, treatments were removed, and the monolayers rinsed once with warm PBS before returning to for a 24-hour recovery period.

For molecular permeability, calcein was applied at 0.5 mM into the apical side of the monolayers with the fruit treatments. After one hour, media in the basal chambers was sampled and examined for fluorescence at 495/515 nm using the plate reader. Application of calibration curves yielded the amount of marker transferred across each monolayer, which was used in the permeability equation $P_{app}=\frac{\Delta M}{C_{a}A\Delta t}$, where P_app_ is the apparent permeability through the monolayer, ΔM is the amount of calcein in the basal compartment, C_a_ is the apical calcein concentration, A is the monolayer area, and Δt is the time between samples. Permeability measurements are expressed as the ratio of each monolayer’s permeability at 3 hours of treatment to its permeability before treatment, normalized to any change in untreated control monolayers during that time.

Cell Signaling Inhibition
Caco-2 monolayers were incubated for an hour before treatment with small molecule inhibitors, then pelargonidin was added without changing the inhibitor concentration. All changes in permeability were normalized to monolayers treated with the inhibitors but no pelargonidin. The inhibitors used were 10 µM FRAX 486 (p21-Activated Kinase, PAK, inhibitor), 10 µM U-73122 (Phospholipase C, PLC, inhibitor) 1 µM SR 3677 (Rho Kinase, ROCK, inhibitor), 1 µM PP2 (c-Src inhibitor), 100 µM API-1 (Protein Kinase B, Akt, inhibitor), 2 µM AZ 628 (Rapidly Accelerated Fibrosarcoma Kinase, Raf, inhibitor), or 0.33 mM PIK (Myosin Light Chain Kinase, MLCK, inhibitor).

Amberlite™ Separation of Strawberry Extracts
 Polyphenols were isolated via a previously published method (*2*). Briefly, a strawberry extract produced by extracting lyophilized fruit with ethanol, then drying via rotary evaporation and lyophilization. The material was dissolved in methanol and adsorbed onto Amberlite™ XAD 7 HP (acrylate ester) resin. The methanol was removed from the resin and evaporated to dryness, yielding unabsorbed material, which comprises a wide variety of compounds. The beads were then washed with water, which was collected and lyophilized to produce a sample composed primarily of sugars and organic acids. Next, the beads were washed with ethanol to collect the remainder of the adsorbed material. The ethanol was removed via rotary evaporation, and any remaining water was lyophilized away to yield a solid, powdered polyphenol extract.

Chromatography
 Medium pressure liquid chromatography (MPLC) was performed using a Buchi Sepacore® system. Glass columns were hand-packed with reverse-phase (C18) silica gel and each run utilized a gradient from 10% - 100% acetonitrile in water with 0.1% trifluoroacetic acid (TFA). Run α was implemented for a coarse separation of the strawberry polyphenol extract (**Table S2**). Eluent absorption at 280 nm was monitored to track phenol group migration. Fractions were collected, concentrated via rotary evaporation, and re-applied to a longer, narrower column for runs β, γ, δ, and ε. Fractions were collected and re-concentrated for testing in cell culture.

Each MPLC fraction was analyzed by ultra performance liquid chromatography (UPLC) using a Waters Acquity UPLC® system and Acquity UPLC C18 Column. Each run began with a 10 μL injection of concentrated MPLC eluent. The samples were separated with a linear gradient of 10% - 100% acetonitrile in water/0.1% TFA over six minutes. (**Table S2**). The eluent was monitored by a photodiode array (PDA) detector, allowing each sample to be recorded for both the 280 nm absorbance trace over time and absorbance spectra of characteristic polyphenol peaks.

To obtain a larger quantity of Fraction ε3 for characterization, a larger amount of strawberry polyphenol material (6.4 g) was used. This material was dissolved in 60 mL of [67% ethanol, 33% water with 0.1% TFA], sonicated for 5 minutes, and centrifuged at 300 x g for 4 minutes to remove any undissolved matter. The supernatant was reduced to 20 mL by rotary evaporation and applied to a 36 x 920 mm column hand-packed with reverse-phase (C18) silica gel. This column was run with the same solvent gradient as Run α, at 50 mL/minute. The corresponding fractions were confirmed via UPLC, concentrated, and subjected to the same conditions as the initial Run ε, yielding approximately 7 mg of sample. Before characterizing, the activity of the sample was confirmed on Caco-2 monolayers (**Fig S6**).

Mass Spectrometry
High-Resolution Mass spectra were obtained on a Thermo Scientific Exactive Plus EMR Orbitrap Mass Spectrometer (ThermoFisher Scientific). The electrospray ionization source (ESI) was operated in the positive mode with spray voltage of 2.5 kV and ion transfer tube temperature at 270°C. Gas flow was set to defaults for the used 10 µL/min injection which was performed using the syringe pump. Each scan consisted of 3 microscans with a detection range set over a mass range of 150–2,000 m/z.

Nuclear Magnetic Resonance (NMR)
 ^1^H and ^13^C NMR spectra were recorded with a Bruker Avance 500 spectrometer (Billerica, MA, USA) using 3 mm tubes with DMSO-*d*_6_ as solvent and tetramethylsilane (TMS) as an internal standard. ^1^H and ^13^C NMR were performed at 500 and 126 MHz, respectively, and chemical shifts are given as δ values.

Mouse Studies
All mouse experiments were approved by the institutional animal care and use committee (IACUC) at Carnegie Mellon University (Pittsburgh, PA, USA) under protocol number PROTO201600017, and were performed in accordance with all institutional, local, and federal regulations. C57BL/6 mice were either purchased from Charles River Laboratories (Wilmington, MA, USA) or obtained from an institutionally managed breeding colony. Prior to experiments, mice were housed in cages of no more than six animals, with controlled temperature (25°C), 12 hour light-dark cycles, and free access to food and water. Mice utilized in this study were female and 8-16 weeks (dextran, intestinal insulin, and toxicity: 18-24 g weight range) or 24–30 weeks old (protein drug capsules: 30–45 g weight range to ensure capsule passage through the gastrointestinal tract). Only mice within 6 weeks of age were directly compared to one another (placed on the same graph) for consistency. The free-to-use PS power calculator (Vanderbilt) was used to determine the minimal sample size for which statistical power was greater than or equal to 0.8. (Generally, n = 5-6). Mice were fasted 8-12 hours the night before an experiment to limit the variability caused by food matter and feces in the GI tract. Fasting also served to stabilize the animals’ blood sugar for insulin activity experiments, with a starting blood glucose range of approximately 70 to 120 mg/dL. Oral gavages were administered at a volume of 10 mL solution per kg of mouse body weight (10 μL/g). Intestinal and subcutaneous injections were administered at a volume of 1 mL/kg (1 μL/g).

Intestinal Permeability to Dextrans
For dextran efficacy studies, fasted mice were orally gavaged with treatment solutions (600 mg/kg STRB PPh or 40 mg/kg pelargonidin), then gavaged one hour later with 600 mg/kg FITC-DX4. Three hours after the dextran gavage, blood was collected and centrifuged. The serum was removed and examined for FITC concentration by reading for fluorescence on the plate reader and comparing to a unique calibration curve for each experiment. For larger macromolecule studies, 40 kDa dextran (FITC-DX40), 70 kDa dextran (FITC-DX70), or 150 kDa dextran (FITC-DX150) was substituted at the same 600 mg/kg dose.

Intestinal Insulin Delivery
 Following ten hours of fasting, mice were orally gavaged with PBS (for control) or strawberry treatments (600 mg/kg STRB PPh or 40 mg/kg pelargonidin). One hour later, their initial blood sugar was measured and the animals were placed under anesthesia. Their intestines were surgically exposed, and insulin was injected at the predetermined dose (1 unit insulin per kg body weight, unless otherwise specified) into the duodenum. The mice were closed and secured with tissue adhesive, then kept under anesthesia as their blood sugar levels were monitored each hour for five hours. An endpoint at five hours was enforced for all experiments, as the combined effects of the anesthesia, dehydration, and reduced blood sugar prevented reliable survival beyond that point. For comparison to the current standard of insulin delivery, subcutaneous injections were given at 1 U/kg to additional mice, into the scruff on their necks. To determine areas above the curve for each mouse, trapezoidal integration was used to sum the area between known points on the blood glucose curve and the starting blood glucose value for the individual animal.

Capsule Preparation
 Dry capsule contents were produced by combining insulin at the prescribed dose, the protease inhibitor aprotinin (25% capsule filling by mass, 25 mg/kg), pelargonidin if required (40% capsule filling by mass, 40 mg/kg), and inactive bovine serum albumin (BSA) filler, then mixing thoroughly. Size M capsules were filled with 3-4 mg of the drug mixture, and their exact weights recorded. Each capsule was then dip coated 3 times in a 7% (w/v in ethanol) solution of Eudragit® L100-55, drying completely under gentle airflow following each coat. The total dry weight of polymer added to each capsule ranged from 0.4 to 0.8 mg.

Oral Insulin Delivery with Capsules

Following a ten-hour fasting period, large (> 30 g) mice were injected subcutaneously with 5 mg/kg metoclopramide hydrochloride (to stimulate gastric emptying) and orally administered capsules. Capsules were chosen so small variations in filler weight matched small variations in mouse weight, giving insulin doses within 10% of the designated average dose. The capsules were immediately flushed into the stomach with a 200 μL gavage of PBS. Blood glucose was measured every two hours for a total of ten hours and normalized to each mouse’s reading before capsule administration. From the blood glucose measurements, areas above the curve were calculated as previously described. These areas were used to calculate dose-corrected relative bioactivity as follows:

$$\begin{matrix} Relative \\ Bioactivity \end{matrix}=\left( \frac{\beta\frac{U}{kg} AAC - 0 \frac{U}{kg} Capsule AAC}{\beta\frac{U}{kg}} \right)/{\left( \frac{1 \frac{U}{kg} SQ AAC-0 \frac{U}{kg} SQ AAC}{1\frac{U}{kg}} \right)\times100\%}$$

where *β* is the insulin dose in U/kg of the capsule treatment being examined, and SQ is subcutaneous injection.

Confocal Microscopy
Monolayers were rinsed with PBS to remove treatments and fixed in 4% paraformaldehyde. They were permeabilized with 0.2% Triton-X100 and blocked with 0.2% BSA solution to limit non-specific antibody binding, then incubated for one hour with staining solutions. The staining solution contained DAPI (12 μg/mL, 358 nm/461 nm) to mark nucleic acids, AlexaFluor 488® conjugated phalloidin (5 units/mL, 495 nm/518 nm) to bind actin, and AlexaFluor® 594 conjugated Anti-ZO-1 antibodies or Anti-Occludin antibodies (50 μg/mL, 590 nm/617 nm) in 0.2% BSA. After staining, the monolayers were mounted on slides using ClearMount™ solution (Invitrogen - Carlsbad, CA, USA) and sealed under coverslips using clear nail polish.

Prepared slides were imaged at 63x magnification using a Zeiss LSM 700 confocal microscope with ZEN 2012 SP1 software. Images were captured using a Plan-Apochromat 63x/1.40 Oil DIC objective and an X-Cite Series 120Q laser source exposing at 405, 488, and 555 nm. Images were approximately 101.5 μm x 101.5 μm and were captured with a lateral resolution of approximately 0.3 μm. No additional processing or averaging was performed to enhance the resolution of the images.

Image J (NIH) image processing software was used to prepare confocal images for publication. Upper and lower thresholds were narrowed slightly to remove background noise and improve visibility of the signals. All images were processed with the same thresholds and display lookup tables (LUTs), which were linear throughout their ranges. Images were converted from their original 16-bit format to RGB color for saving and arrangement into figures. No other manipulations were performed.

Long-Term Safety Study

Cohorts of 12 mice each were treated for thirty days with oral pelargonidin (40 mg/kg) or PBS (negative control). Mice were weighed each day before oral gavages to ensure proper dosing. On day 31, half of the mice were sacrificed, with tissue and serum collected for later analysis. The remaining six mice per group were left untreated for an additional seven days to assess their recovery from any cumulative effects of the pelargonidin treatment. Reported weights are normalized to the starting weight of the individual mouse, and to the untreated control mice.

To determine levels of circulating cytokines and inflammatory markers, blood samples were separated via centrifugation. The serum was subjected to ELISA analysis per the instructions of the kit manufacturer.

For intestinal tight junction expression, tissue samples (1 cm) were collected from treated mice and homogenized in 0.5 mL Trizol reagent using the BeadBug microtube homogenizer. Chloroform (0.1 ml) was added and centrifuged at 12,000 rpm for 15 min. The aqueous layer was removed and mixed with an equal volume of 100% ethanol, and the RNA was extracted with RNeasy mini kit (Qiagen) according to the manufacturer’s instructions. cDNA was generated from 2,000 ng of RNA using High-Capacity cDNA Reverse Transcription Kit (Applied Biosystems). Quantitative real-time polymerase chain reaction (qRT-PCR) was performed using SYBR Select Master Mix (Applied Biosystems) on a ViiA 7 Real-Time PCR system (Applied Biosystems) and the primer sequences displayed in **Table S3**. The expression of each mRNA was normalized to expression of the housekeeping gene Beta Actin and expressed as R=2^-ΔΔct values.

Histology

Immediately following euthanasia, mice from the safety study were dissected to collect their small intestines and colons. The organs were fixed in 4% formaldehyde for 24 hours and transferred into 30% sucrose solution overnight at 4°C. Sampled were embedded in OCT, and cross-sections of 10 µm were cut using a cryostat. Sections were rinsed with PBS and stained in Harris hematoxylin for eight minutes, then washed twice in DI water, differentiated for one minute in 5% acetic acid, and washed with water two more times. The hematoxylin was blued using Scott’s tap water (0.2 % sodium bicarbonate and 1% magnesium sulfate at pH 8 in water) and washed twice more with water. The slides were stained in alcoholic eosin for two minutes, then dehydrated with 95% and 100% ethanol before they were cleared in xylenes. The slides were mounted and allowed to dry in a fume hood overnight, then imaged using an EVOS FL Auto 2 microscope (Thermo Fisher Scientific) at 20x magnification.

Statistics
 All data presented as arithmetic mean of the given “n” number of biological replicates (individual animals or number of in vitro cell culture wells), and error bars display the standard error of the mean. For statistical significance markings, two-tailed Student’s t-tests with Welch’s correction were used to calculate p values, which are displayed alongside the relevant data.

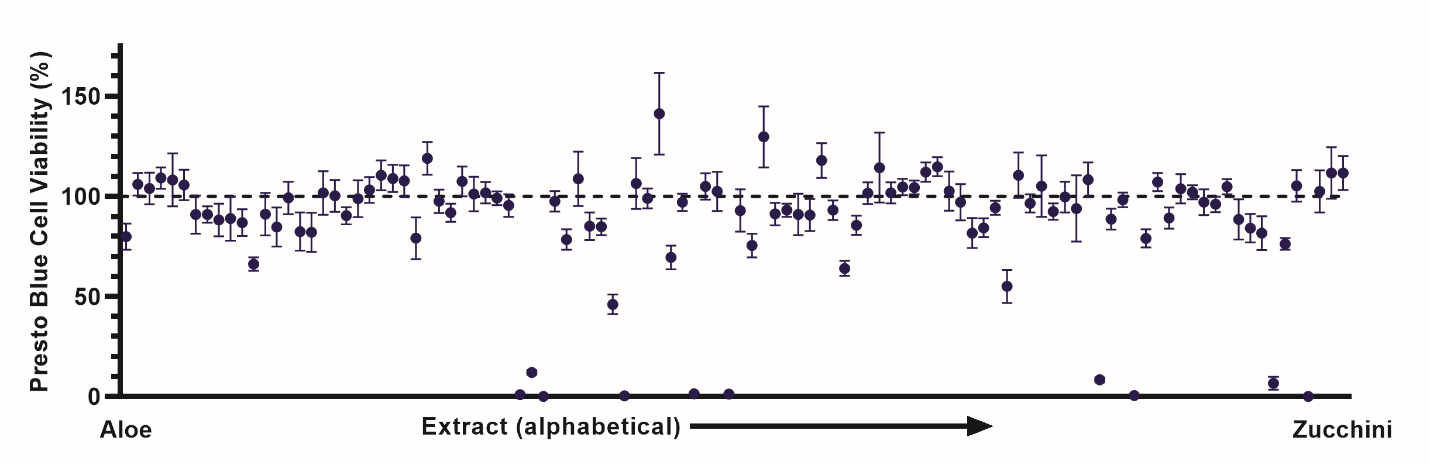

Figure S1. Crude extracts were generally well-tolerated by Caco-2 cells. By the PrestoBlue® viability assay, only a handful of extracts were toxic to intestinal cells following three hours of exposure. Error bars display s.e.m. (n = 4).

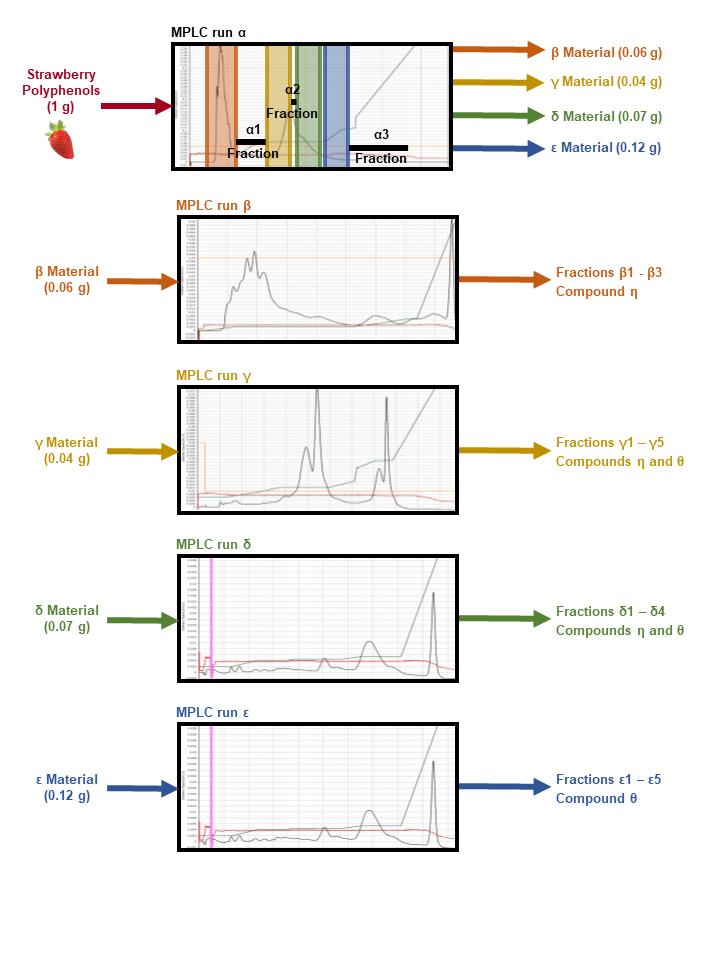

**Figure S2: An MPLC separation scheme was used to resolve the strawberry polyphenols into 22 fractions.** Each MPLC run was assigned a Greek letter identifier, moving consecutively through the alphabet. The first MPLC run (α) yielded fractions that were divided into seven groups, based on fraction color and ultra performance liquid chromatography (UPLC) traces of the eluents. Four of these groups produced enough material to support a second tier of MPLC runs, yielding fractions for the β, γ, δ, and ε runs, respectively. UPLC traces identified two pure compounds, each of which appeared as discrete fractions in three of the four runs. Fractions with identical HPLC traces of a single compound were pooled together to yield the samples denoted as Compound η and Compound θ.

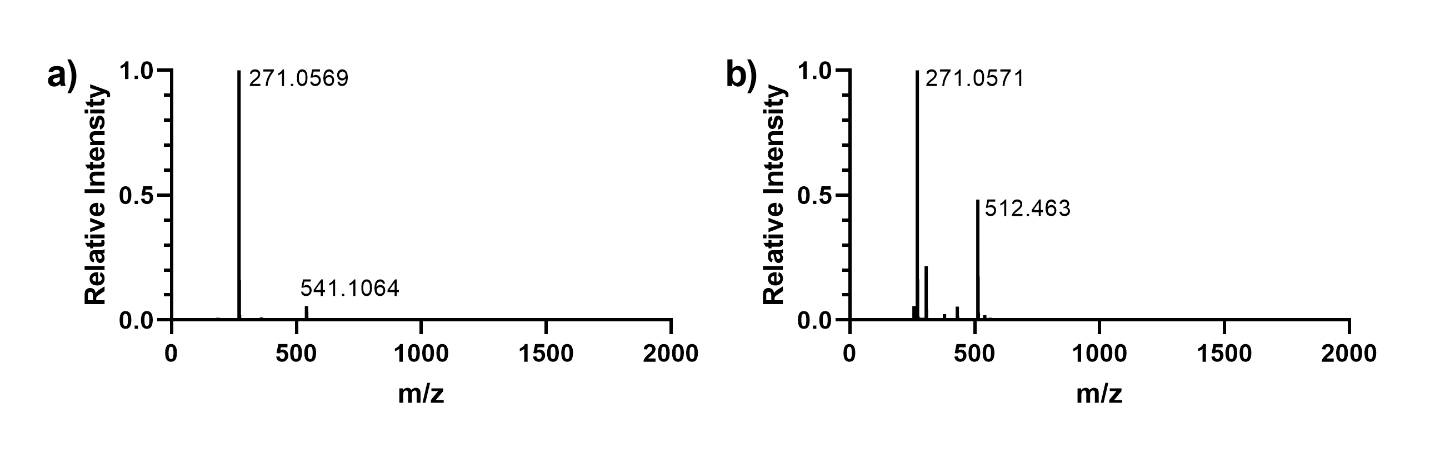

**Figure S3: ESI-TOF mass spectrometry confirmed the active component in strawberry to be pelargonidin. (a)** While both commercially purchased pelargonidin and **(b)** the MPLC fraction ε3 contained a small number of impurities, the peak of interest for protonated pelargonidin (271 g/mol), in positive ion mode, was the most prominent in each case.

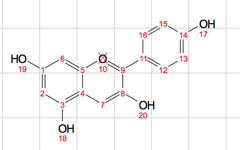

**Assignments:**

#1	167.79

#2	6.88/102.64

#3	155.68

#4	112.83

#5	157.10

#6	6.97/95.27

#7	8.73/134.15

#8	160.74

#9	145.64

#11	120.45

#12,16	8.53/133.75

#13,15	7.10/117.17

#14	164.51

**Figure S4: NMR confirmed that fraction ε3 was the same chemical entity as commercially purchased pelargonidin. Standard ^1^H NMR** (500 MHz, DMSO-*d*_6_) δ 12.16 (m, 3H), 11.27 (s, 1H), 8.86 – 8.70 (m, 1H), 8.55 (dd, *J* = 9.0, 1.8 Hz, 2H), 7.25 – 7.06 (m, 2H), 6.96 (d, *J* = 1.9 Hz, 1H), 6.87 – 6.77 (m, 1H). **Standard ^13^C NMR** (126 MHz, DMSO) δ 167.64, 164.52, 160.06, 157.01, 155.70, 145.59, 134.04, 133.87, 120.45, 117.17, 112.80, 102.57, 94.34. **Fraction ^1^H NMR** (500 MHz, DMSO-*d*_6_) δ 12.31 (d, *J* = 80.7 Hz, 3H), 11.36 (s, 1H), 8.96 – 8.82 (m, 1H), 8.53 (dd, *J* = 9.3, 2.4 Hz, 2H), 7.20 – 7.07 (m, 2H), 6.99 (t, *J* = 2.1 Hz, 1H), 6.88 (dt, *J* = 5.9, 2.6 Hz, 1H). **Fraction ^13^C NMR** (126 MHz, DMSO) δ 167.79, 164.51, 160.74, 157.10, 155.68, 145.64, 134.15, 133.75, 120.44, 117.17, 112.83, 102.64, 94.27. Assignments are given based on the NMR readout from the fraction run.

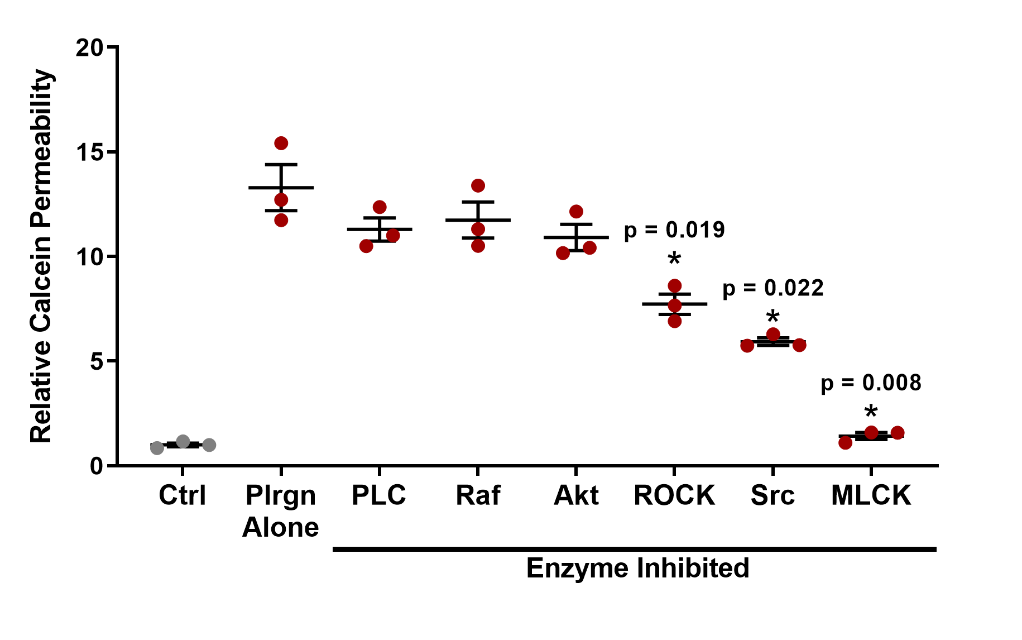

**Figure S5: Pelargonidin increases epithelial permeability by an MLCK-dependent mechanism.** Pre-treatment of Caco-2 intestinal cells with small molecule inhibitors of several epithelial signaling enzymes did not affect the efficacy of pelargonidin. However, the presence of an MLCK inhibitor completely eliminated any permeation enhancing effects. Src and ROCK may be partially involved in the signaling that directs opening of tight junctions, but there are likely more pathways between pelargonidin and MLCK than only these two. Error bars display s.e.m. (n = 3). * p < 0.05 by Student’s t-test with Welch’s correction.

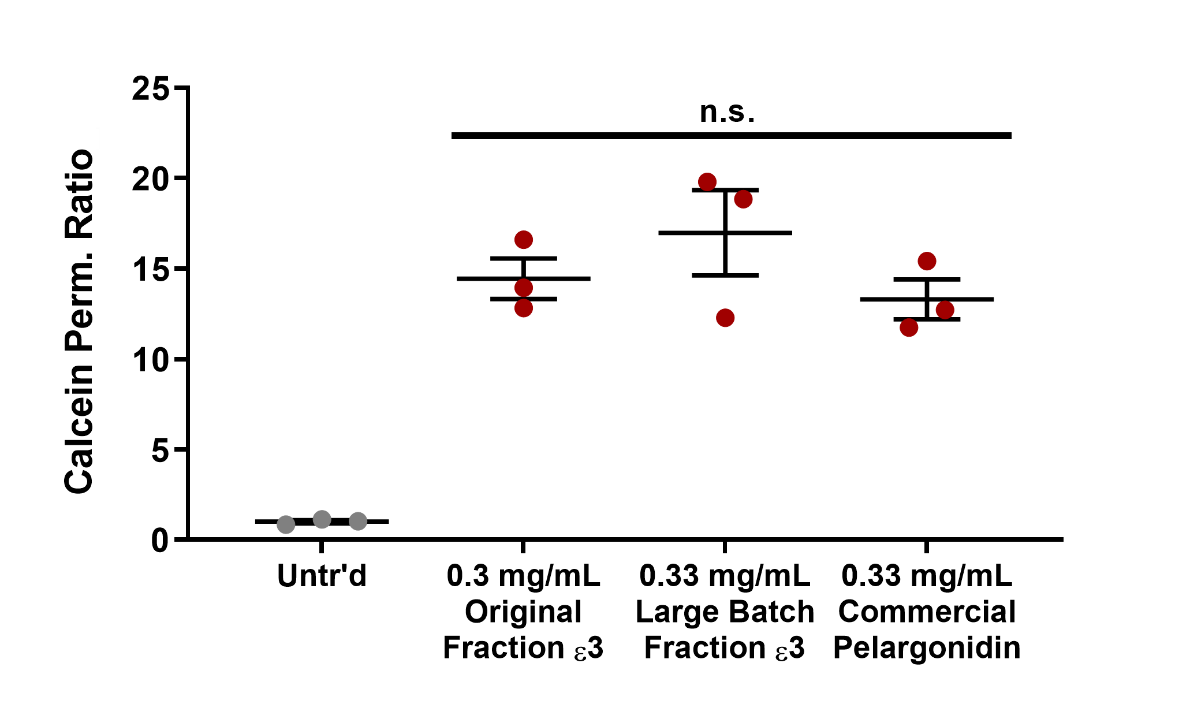

**Figure S6: Similar permeation enhancing activity was confirmed for active chromatography fractions and pelargonidin reference standard.** There were no statistically significant differences in the calcein permeability ratio between the commercially obtained pelargonidin (reference standard) and the two batches of active chromatography fractions ε3. through treated Caco-2 monolayers. Error bars display s.e.m. (n = 3); n.s. = not statistically significant by Student’s t-test with Welch’s correction.

Table S1. Presto viability, TEER, and calcein permeability screening results for crude food extracts. Viability (“Via.”) columns are all presented with respect to untreated control cells. Some of the toxic extracts were not tested for TEER or calcein permeability. Lower values for "TEER % 3 h" denote more efficacious permeation enhancers. "TEER Rec." is short for TEER Recovery and denotes whether the average TEER value for the treated monolayers returned to at least 80% of its original value 24 hours after extract removal. "Calcein Class" indicates whether the extract (P)ermeabilized the monolayers or (C)losed tight junctions to a statistically significant extent, or caused (I)nsignificant differences from control. * p < 0.05 by Student's two-tailed t-test with respect to untreated control.

| **Extract** | **Via. %** | **Via. s.e.m.** | ***** |  | **TEER % 3 h** | **TEER s.e.m.** | **TEER Rec.** | **Calcein Ratio** | **Calcein s.e.m.** | **Calcein Class** |
| --- | --- | --- | --- | --- | --- | --- | --- | --- | --- | --- |
| Aloe | 79.9 | 6.6 | Yes |  | 2.6 | 2.7 | No | 28.6 | 3.2 | P |
| Apple, Granny Smith | 106.0 | 5.6 | No |  | 73.1 | 6.9 | Yes | 1.3 | 0.2 | I |
| Apple, Red Delicious | 103.9 | 7.9 | No |  | 110.3 | 13.4 | Yes | 0.8 | 0.1 | I |
| Asparagus | 109.1 | 5.3 | No |  | 30.5 | 9.7 | No | 1.8 | 1.1 | I |
| Avocado | 108.2 | 13.2 | No |  | 98.4 | 8.5 | Yes | 1.5 | 0.4 | I |
| Banana | 105.6 | 7.6 | No |  | 90.0 | 3.7 | Yes | 0.7 | 0.1 | I |
| Basil | 90.9 | 9.6 | No |  | 13.4 | 1.3 | No | 9.5 | 2.3 | P |
| Beans, Green | 90.9 | 4.0 | No |  | 51.6 | 6.4 | Yes | 1.3 | 0.3 | I |
| Beet | 88.2 | 8.2 | No |  | 2.8 | 0.6 | No | 52.5 | 18.2 | P |
| Blackberry | 88.9 | 11.0 | No |  | 106.9 | 8.5 | Yes | 0.8 | 0.1 | I |
| Blueberry | 86.9 | 6.8 | No |  | 93.9 | 6.2 | Yes | 0.5 | 0.2 | I |
| Breadfruit | 66.2 | 3.4 | Yes |  |  |  |  |  |  |  |
| Broccoli | 91.0 | 10.7 | No |  | 76.8 | 12.0 | Yes | 0.7 | 0.0 | C |
| Brussels Sprouts | 84.6 | 9.8 | No |  | 90.8 | 4.9 | Yes | 0.8 | 0.1 | I |
| Cabbage, Red | 99.2 | 8.1 | No |  | 73.6 | 10.9 | Yes | 1.1 | 0.2 | I |
| Cactus Pear | 82.4 | 9.6 | No |  | 101.9 | 1.8 | Yes | 0.7 | 0.1 | I |
| Cantaloupe | 82.0 | 9.7 | No |  | 106.5 | 8.5 | Yes | 1.0 | 0.2 | I |
| Carrots, Orange | 101.7 | 10.9 | No |  | 97.6 | 3.1 | Yes | 1.4 | 0.7 | I |
| Carrots, Purple | 100.2 | 7.9 | No |  | 115.4 | 11.5 | Yes | 1.8 | 1.2 | I |
| Celery | 90.3 | 4.3 | No |  | 87.0 | 7.4 | Yes | 1.0 | 0.1 | I |
| Chayote | 98.8 | 9.2 | No |  | 20.1 | 1.6 | Yes | 4.7 | 0.6 | P |
| **Table S1, continued** | | | | | | | | | | |
| **Extract** | **Via. %** | **Via. s.e.m.** | ***** |  | **TEER % 3 h** | **TEER s.e.m.** | **TEER Rec.?** | **Calcein Ratio** | **Calcein s.e.m.** | **Calcein Class** |
| Cherry, Red | 103.2 | 6.4 | No |  | 95.4 | 1.7 | Yes | 0.9 | 0.1 | I |
| Cherry, White | 110.4 | 7.4 | No |  | 95.6 | 1.3 | Yes | 0.9 | 0.1 | I |
| Chokeberry | 108.9 | 6.8 | No |  | 32.9 | 2.4 | No | 0.8 | 0.1 | I |
| Cilantro | 107.7 | 7.7 | No |  | 60.9 | 11.5 | No | 0.9 | 0.1 | I |
| Corn | 79.1 | 10.5 | No |  | 99.5 | 25.2 | Yes | 1.1 | 0.1 | I |
| Cranberry | 118.9 | 8.2 | No |  | 16.3 | 1.6 | No | 5.2 | 0.3 | P |
| Cucumber | 97.5 | 5.8 | No |  | 54.7 | 14.8 | Yes | 1.7 | 0.3 | I |
| Currant, Black | 91.8 | 4.5 | No |  | 22.6 | 2.5 | No | 1.9 | 0.3 | I |
| Currant, Red | 107.4 | 7.4 | No |  | 20.3 | 0.5 | No | 2.6 | 0.3 | P |
| Dill | 101.1 | 8.6 | No |  | 43.7 | 6.2 | No | 1.6 | 0.1 | P |
| Dragonfruit | 101.8 | 5.3 | No |  | 87.5 | 11.6 | Yes | 0.9 | 0.1 | I |
| Eggplant | 99.1 | 3.4 | No |  | 92.1 | 6.7 | Yes | 0.9 | 0.1 | I |
| Fennel | 95.4 | 5.6 | No |  | 36.2 | 10.8 | Yes | 2.4 | 0.8 | P |
| Galangal | 1.0 | 1.4 | Yes |  |  |  |  |  |  |  |
| Garlic | 12.0 | 1.2 | Yes |  | 18.5 | 3.3 | No | 5.5 | 1.1 | I |
| Ginger | -0.1 | -0.8 | Yes |  | 1.6 | 1.1 | No | 125.6 | 59.3 | I |
| Grapefruit | 97.5 | 5.1 | No |  | 95.3 | 10.1 | Yes | 0.9 | 0.1 | I |
| Grapefruit Rind | 78.4 | 5.1 | Yes |  |  |  |  |  |  |  |
| Grapes, Green | 108.8 | 13.6 | No |  | 70.0 | 2.9 | Yes | 0.9 | 0.2 | I |
| Grapes, Red Seedless | 85.0 | 6.9 | No |  | 16.7 | 0.6 | Yes | 6.2 | 0.7 | P |
| Guava | 84.8 | 4.2 | Yes |  |  |  |  |  |  |  |
| Horseradish | 46.1 | 4.9 | Yes |  |  |  |  |  |  |  |
| Huckleberry, Garden | 0.4 | 0.2 | Yes |  |  |  |  |  |  |  |
| Jalapeno | 106.4 | 12.6 | No |  | 57.7 | 12.6 | Yes | 1.7 | 0.6 | I |
| Jaltomato | 98.9 | 4.9 | No |  | 42.2 | 0.6 | Yes | 1.7 | 0.1 | P |
| Jicama | 141.2 | 20.5 | No |  | 91.7 | 5.8 | No | 0.9 | 0.1 | I |
| Kiwi | 69.4 | 5.9 | Yes |  |  |  |  |  |  |  |
| Lemon | 97.0 | 4.3 | No |  | 14.7 | 0.9 | No | 6.8 | 0.7 | P |
| **Table S1, continued** | | | | | | | | | | |
| **Extract** | **Via. %** | **Via. s.e.m.** | ***** |  | **TEER % 3 h** | **TEER s.e.m.** | **TEER Rec.?** | **Calcein Ratio** | **Calcein s.e.m.** | **Calcein Class** |
| Lemon Rind | 1.3 | 0.2 | Yes |  |  |  |  |  |  |  |
| Lettuce | 104.9 | 6.6 | No |  | 65.1 | 7.8 | Yes | 2.6 | 0.9 | I |
| Lime | 102.4 | 9.8 | No |  | 78.0 | 16.0 | Yes | 2.7 | 0.8 | P |
| Lime Rind | 1.2 | 0.2 | Yes |  |  |  |  |  |  |  |
| Mango | 92.9 | 10.6 | No |  | 101.5 | 4.2 | Yes | 1.7 | 0.2 | I |
| Mushroom | 75.4 | 5.9 | Yes |  | 42.6 | 2.4 | No | 2.5 | 0.5 | I |
| Name | 129.7 | 15.3 | No |  | 68.9 | 14.6 | Yes | 1.0 | 0.6 | I |
| Nectarine | 91.2 | 5.6 | No |  | 57.7 | 5.0 | Yes | 3.0 | 0.7 | P |
| Okra | 93.1 | 3.3 | No |  | 120.7 | 20.2 | Yes | 0.8 | 0.1 | I |
| Onion, Red | 90.9 | 10.3 | No |  | 75.2 | 7.8 | Yes | 1.0 | 0.1 | I |
| Onion, Yellow | 90.6 | 8.0 | No |  | 71.7 | 6.3 | Yes | 1.3 | 0.1 | I |
| Orange | 117.9 | 8.7 | No |  | 108.7 | 5.0 | Yes | 0.5 | 0.2 | I |
| Orange Rind | 64.0 | 3.7 | Yes |  |  |  |  |  |  |  |
| Otricoli Orange Berry | 93.1 | 4.9 | No |  | 48.3 | 3.8 | Yes | 1.2 | 0.1 | I |
| Papaya | 85.5 | 4.8 | Yes |  |  |  |  |  |  |  |
| Parsley | 101.5 | 5.4 | No |  | 30.7 | 8.6 | Yes | 5.0 | 0.7 | P |
| Parsnip | 114.3 | 17.5 | No |  | 95.7 | 13.6 | Yes | 1.1 | 0.1 | I |
| Passionfruit | 101.8 | 5.3 | No |  | 41.9 | 7.3 | Yes | 2.4 | 0.3 | P |
| Peach | 104.7 | 4.1 | No |  | 111.4 | 10.6 | Yes | 0.3 | 0.2 | I |
| Peach, White | 104.4 | 3.4 | No |  | 120.8 | 9.2 | Yes | 0.5 | 0.3 | I |
| Pear, Anjou | 112.1 | 4.9 | No |  | 90.5 | 9.3 | Yes | 2.7 | 0.4 | I |
| Pear, Asian | 114.8 | 4.8 | No |  | 87.7 | 20.0 | Yes | 3.9 | 2.1 | I |
| Pepino | 102.6 | 9.7 | No |  | 29.7 | 3.3 | Yes | 2.3 | 0.2 | P |
| Pepper, Green Bell | 97.1 | 9.1 | No |  | 43.5 | 3.5 | No | 1.9 | 0.2 | P |
| Pepper, Red Bell | 81.7 | 7.5 | Yes |  | 88.5 | 1.8 | Yes | 0.9 | 0.2 | I |
| Peppermint | 84.2 | 4.8 | No |  | 41.3 | 11.3 | Yes | 3.6 | 0.9 | P |
| Persimmon | 94.3 | 3.5 | No |  | 89.6 | 14.2 | Yes | 1.6 | 0.7 | I |
| Pineapple | 55.0 | 8.3 | Yes |  |  |  |  |  |  |  |
| **Table S1, continued** | | | | | | | | | | |
| **Extract** | **Via. %** | **Via. s.e.m.** | ***** |  | **TEER % 3 h** | **TEER s.e.m.** | **TEER Rec.?** | **Calcein Ratio** | **Calcein s.e.m.** | **Calcein Class** |
| Plum, Purple | 110.5 | 11.3 | No |  | 40.9 | 3.5 | Yes | 1.6 | 0.2 | I |
| Plum, Yellow | 96.5 | 4.5 | No |  | 63.4 | 3.6 | Yes | 1.1 | 0.1 | I |
| Pomegranate | 105.1 | 15.4 | No |  | 104.1 | 9.1 | Yes | 0.9 | 0.1 | I |
| Pomelo | 92.4 | 4.1 | No |  | 72.4 | 14.3 | Yes | 0.8 | 0.4 | I |
| Pomelo Rind | 99.6 | 7.7 | No |  | 61.8 | 6.9 | Yes | 0.8 | 0.4 | I |
| Potato, Baking | 93.9 | 16.6 | No |  | 36.7 | 2.9 | Yes | 4.4 | 2.0 | I |
| Potato, Blue | 108.2 | 8.7 | No |  | 30.3 | 3.0 | Yes | 5.2 | 2.2 | I |
| Potato, Red | 8.3 | 1.3 | Yes |  | 3.7 | 0.6 | No | 28.4 | 2.5 | P |
| Quince | 88.6 | 5.3 | No |  | 75.9 | 15.2 | No | 1.5 | 0.9 | I |
| Raspberry | 98.2 | 3.6 | No |  | 120.6 | 5.6 | Yes | 0.3 | 0.1 | C |
| Rosemary | 0.6 | 0.3 | Yes |  |  |  |  |  |  |  |
| Spinach | 79.0 | 4.6 | Yes |  |  |  |  |  |  |  |
| Starfruit | 107.1 | 4.6 | No |  | 13.7 | 1.5 | Yes | 5.1 | 0.6 | P |
| Strawberry | 89.1 | 5.3 | No |  | 17.8 | 2.9 | Yes | 7.9 | 0.5 | P |
| Strawberry, White | 103.8 | 7.3 | No |  | 60.6 | 17.3 | Yes | 1.6 | 0.2 | P |
| Sugar Cane | 102.0 | 3.5 | No |  | 101.8 | 11.1 | Yes | 4.1 | 3.3 | I |
| Sweet Potato | 97.0 | 6.5 | No |  | 99.3 | 4.7 | Yes | 1.0 | 0.4 | I |
| Sweet Potato, Purple | 96.0 | 3.8 | No |  | 78.0 | 5.4 | No | 1.2 | 0.2 | I |
| Thyme | 104.8 | 3.8 | No |  | 29.5 | 4.4 | Yes | 1.8 | 0.1 | P |
| Tomato, Black | 88.4 | 10.0 | No |  | 100.5 | 5.9 | Yes | 0.5 | 0.1 | C |
| Tomato, Red | 84.1 | 7.2 | No |  | 89.3 | 10.2 | Yes | 3.7 | 0.8 | P |
| Tomato, Yellow | 81.6 | 8.5 | No |  | 102.4 | 7.6 | Yes | 1.0 | 0.2 | I |
| Turmeric | 6.7 | 3.2 | Yes |  |  |  |  |  |  |  |
| Turnip | 76.2 | 2.9 | Yes |  |  |  |  |  |  |  |
| Watermelon | 105.3 | 7.9 | No |  | 117.6 | 8.0 | Yes | 0.5 | 0.2 | I |
| Wonderberry | 0.0 | 0.5 | Yes |  |  |  |  |  |  |  |
| Yautia | 102.4 | 10.5 | No |  | 91.8 | 10.8 | No | 0.7 | 0.1 | C |
| Yucca | 111.7 | 12.8 | No |  | 104.7 | 12.5 | Yes | 0.9 | 0.5 | I |
| Zucchini | 111.7 | 8.5 | No |  | 81.8 | 8.1 | Yes | 0.8 | 0.1 | I |

Table S2. Parameters for Chromatography Runs. In each case, solvent A is 0.1 % trifluoroacetic acid (TFA) in water and solvent B is acetonitrile

| **MPLC Run α**  Column: 26 x 460 mm | | |  | **MPLC Runs β, γ, δ, ε**  Column: 15 x 920 mm | | |  | **UPLC**  Acquity UPLC Column | | |
| --- | --- | --- | --- | --- | --- | --- | --- | --- | --- | --- |
| Starting mass: 1 g Pph | | |  | Starting mass: | | |  | Starting material: | | |
| Time (min) | % A | % B |  | β: 0.06 g | | |  | 10 μL MPLC fraction | | |
|  |  |  |  | γ: 0.04 g | | |  | Time (min) | % A | % B |
| 0.00 | 90 | 10 |  | δ: 0.07 g | | |  |  |  |  |
| 4.87 | 90 | 10 |  | ε: 0.12 g | | |  | 0.00 | 90 | 10 |
| 7.57 | 80 | 20 |  | Time (min) | % A | % B |  | 1.00 | 85 | 15 |
| 11.25 | 80 | 20 |  |  |  |  |  | 3.50 | 80 | 20 |
| 13.53 | 70 | 30 |  | 0.00 | 90 | 10 |  | 6.00 | 0 | 100 |
| 17.50 | 70 | 30 |  | 6.05 | 90 | 10 |  | 7.00 | 0 | 100 |
| 28.40 | 60 | 40 |  | 14.93 | 85 | 20 |  |  |  |  |
| 39.12 | 60 | 40 |  | 20.98 | 80 | 20 |  |  |  |  |
| 46.93 | 0 | 100 |  | 28.65 | 0 | 100 |  |  |  |  |
| 51.00 | 0 | 100 |  | 32.31 | 0 | 100 |  |  |  |  |

**Table S3: Primer sequences used for qRT-PCR**

| Gene | Forward | Reverse |
| --- | --- | --- |
| BACTIN | CACTGTCGAGTCGCGTCC | TCATCCATGGCGAACTGGTG |
| CLDN2 | GAAAGGACGGCTCCGTTTTC | CAGTGTCTCTGGCAAGCTGA |
| CLDN5 | GTTAAGGCACGGGTAGCACT | TACTTCTGTGACACCGGCAC |
| ZO1 | CTCTTCAAAGGGAAAACCCGA | GTACTGTGAGGGCAACGGAG |
| JAMA | TCCCGAGAACGAGTCCATCA | GAACTTCCACTCCACTCGGG |
